# Caudal-rostral progression of alpha motoneurone degeneration in the SOD1^G93A^ mouse model of amyotrophic lateral sclerosis

**DOI:** 10.1101/2020.06.27.173054

**Authors:** Alastair J Kirby, Thomas Palmer, Richard Mead, Ronaldo Ichiyama, Samit Chakrabarty

**Author notes:** Corresponding author: Samit Chakrabarty^1^, 0113 3434256.

## Abstract

Mice with transgenic expression of human SOD1^G93A^ are a widely used model of ALS, with a caudal-rostral progression of motor impairment. Previous studies have quantified the progression of motoneurone (MN) degeneration based on size, even though alpha (α-) and gamma (γ-) MNs overlap in size. Therefore, using molecular markers and synaptic inputs, we quantified the survival of α-MNs and γ-MNs at the lumbar and cervical spinal segments of 3- and 4-month SOD1^G93A^ mice, to investigate whether there is a caudal-rostral progression of MN death. By 3-months, in the cervical and lumbar spinal cord, there was α-MN degeneration with complete γ-MN sparing. At 3-months the cervical spinal cord had more α-MNs per ventral horn than the lumbar spinal cord, in SOD1^G93A^ mice. A similar spatial trend of degeneration was observed in the corticospinal tract, which remained intact in the cervical spinal cord at 3- and 4-months of age. These findings agree with the corticofugal synaptopathy model, that α-MN and CST of the lumbar spinal cord are more susceptible to degeneration in SOD1^G93A^ mice. Hence, there is spatial and temporal caudal-rostral progression of α-MN and CST degeneration in SOD1^G93A^ mice.

**Highlights:**

1. SOD1^G93A^ mice display a caudal-rostral progression of motor impairment.
2. Lumbar spinal cord of SOD1^G93A^ mice has an enhanced susceptibility to degeneration.
3. SOD1^G93A^ mice exhibit a caudal-rostral progression of α-MN and CST degeneration

## Introduction

ALS is a rare disorder of the motor system characterised by selective degeneration of the force-producing α-motoneurones (α-MN). Accordingly, in humans, the site of symptomatic onset corresponds to the greatest motor neuron death [1].

Mice with transgenic expression of the human SOD1^G93A^ gene are a widely used model of ALS and symptoms of motor deficits begin with initial decline in rotarod performance, leading to defect of gait [2]. Gait becomes increasingly forelimb dependent, and hindlimb paralysis precedes forelimb paralysis at the end stage [2].

Previous studies have quantified the motoneurone degeneration in the SOD1^G93A^ mouse to correlate motoneurone loss to motor impairment [3]. However, some motoneurone subtypes are less susceptible to degeneration than others. In mouse models of ALS, proprioceptive γ-motoneurones (γ-MN) appear to be spared from degeneration [4,5].

Previous studies were reliant on soma size to differentiate between α- and γ-MNs, which due to an overlap in size of the populations of α- and γ-MNs, is inaccurate. However, α- and γ-MNs can be distinguished from each through the selective down regulation of NeuN and absence of cholinergic input on the soma of γ-MNs [4,6,7].

Using selective molecular markers, we quantified the survival of α- and γ-MNs at the lumbar and cervical spinal segments of 3- and 4-month SOD1^G93A^ mice, to investigate whether there is a caudal-rostral progression of α-MN death, concurrent with motor deficits. By 3-months SOD1^G93A^ mice exhibit visible impairment and tremor of the hindlimb. These symptomatic signs correlate with a reduction in lower limb volume and a decline in rotarod performance. At 4-months of age, SOD1^G93A^ mice demonstrate impairment of gait and rotarod performance with extensive muscle loss [2]. Due to the sharp decline in motor performance between 3- and 4-months, we selected these time points to assess the caudal-rostral progression of α-MN death.

The basis of the deterioration of motor control in ALS is due to dysfunction and degeneration of both upper (supraspinal) and lower (α) motoneurones, but the pathophysiology remains contested. Specifically, whether motoneurone dysfunction begins at the neuromuscular junction and propagates retrogradely (‘dying-back’) or begins at the motor cortex and propagates anterogradely (‘dying-forward’) [8]. The corticofugal synaptopathy model supports evidence of motoneurone death propagating from the cortex but also implicates the cortical-motor synapse in ALS pathology [9]. This model would suggests that longer, lumbar-targeting supraspinal pyramidal neurones are more vulnerable to degeneration, creating a distal to proximal progression of ALS [10,11]. We, therefore, measured the integrity of the corticospinal tract using PKC-γ (Protein kinase C-γ) [12,13] at cervical and lumbar levels.

## Materials and methods

C57BL/6 J hSOD1^G93A^ transgenic mice were as described in [2]. Wild-type (WT) age-matched controls were purchased from Harlan Laboratories. The number and sex of mice used are displayed in supplementary table 1. Experiments were in accordance with the UK animal scientific procedures act of 1986.

Mice were transcardially perfused with 1M PBS followed by 4% paraformaldehyde in PBS, spinal cords were dissected and post fixed with 4% paraformaldehyde/PBS for 24 hours before transferring to PBS. Sections were stored in a sucrose based cryoprotectant at −70°C until sectioning.

### Immunohistochemistry

Twenty-five μm thick coronal sections of the phrenic spinal segments (C3-C5), the lower cervical segments (C6-C8), and the upper lumbar (L1-L4) spinal cord were cut using a cryostat (Leica CM1850). Every 3^rd^/4^th^ sections were taken for staining.

Sections were washed in 0.1M PBS solution before blocking by incubation with 3% normal donkey serum (NDS, Sigma; D9663), 0.2% PBS-Triton for 2 hours. Sections were incubated with goat polyclonal anti-ChAT primary antibodies (1:500, Millipore; AB144P) for 48 hours at 4°C in 3% NDS, 0.2% Triton-x in PBS Following three washes in 0.1M PBS, sections were incubated with mouse monoclonal anti-NeuN primary antibodies (1:500, Millipore; MAB377) for 24 hours at 4°C in 3% NDS, 0.2% PBST. After three 10-minute washes in 1xPBS, sections were incubated with anti-goat Alexa Fluor 488 (1:500, Invitrogen; A11055) and anti-mouse Alexa Fluor 647 (1:500, Invitrogen; A31570) conjugated secondary antibodies in 3% NDS, 0.2% Triton-x in PBS for 2 hours. Following two washes in 0.1 M PBS, the sections were mounted using Vectashield (Vector lab, H-1000).

To visualise the integrity of the corticospinal tract (CST), three lumbar and three cervical sections for each mouse were subject to the same procedure but incubated with rabbit polyclonal anti-PKCγ (1:500, Santa Cruz Biotechnology; sc-211). Anti-rabbit Alexa Fluor 555 conjugated secondary antibodies (1:500, Invitrogen; A31572) were applied as above.

### Imaging and analysis

Imaging of the stained sections was performed by a Zeiss LSM700 inverted confocal microscope, using an 40x objective. Z-stacks of 3μm depth were taken of 8 ventral horns (VH) per animal per spinal cord level, to avoid double counting of cells.

We identified 3 subpopulations of MN in the Lumbar and Cervical sections. α-MNs were distinguished from γ-MNs using the selective expression of NeuN and presence of C boutons on the soma, [6,7]. Two groups of ChAT^+^ α-MNs were identified, α-MNs which were also NeuN positive (ChAT^+^/NeuN^+^/C-bouton^+^) and α_p_-MNs where NeuN is downregulated (ChAT^+^/NeuN^-^/C-bouton^+^). ChAT negative MN were identified as γ-MNs (ChAT^+^/NeuN^-^/C-bouton^-^) (Fig 1).

**Figure 1:**
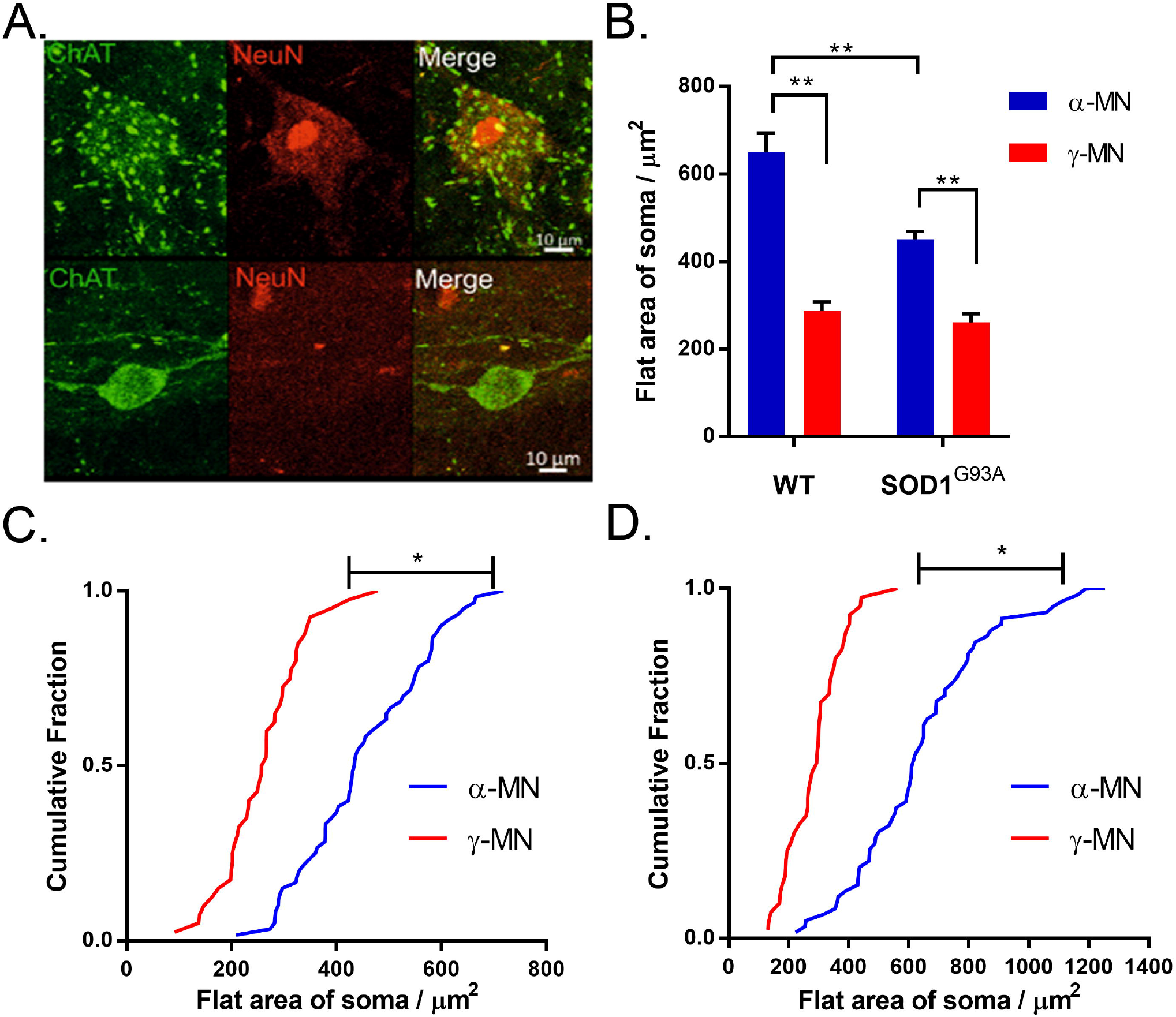
α- and γ-MNs can be distinguished by selective expression of ChAT and NeuN. A) α-MN staining ChAT^+^/NeuN^+^/C-bouton^+^, γ-MN staining ChAT^+^/NeuN^-^/C-bouton^-^. Both images are projections through the z-plane of one section. Scale bar = 10 μm. B) <u>Larger</u> α-MNs lost preferentially in SOD1^G93A^ mice. Significance was determined by a two-way ANOVA and Tukey’s post-hoc test. Columns represent mean ± S.E.M., C) cumulative fraction plot of the size distribution of α-MNs between SOD1^G93A^ mice and wild type controls (N α-MN SOD1^G93A^ (60) WT (60)), *p<0.01 D) cumulative fraction plot of the size distribution of γ-MNs between SOD1^G93A^ mice and wild type controls (N γ-MN: SOD1^G93A^ (40) WT (40)). Significance was determined with a Mann Whitney u test due to non-normal distribution, p>0.05 E) α_p_-MNs staining ChAT^+^/NeuN^+^/C-bouton^+^. F) Cumulative fraction plot of α_p_-MNs and α-MN flat soma area. Mann Whitney u test was used to determine significance, p>0.05.

Image acquisition and counting was performed using Zeiss Zen 2.3 and FIJI [14]. In two mice per group, the soma area of 15 α-MNs and 10 γ-MNs was measured by defining the soma in the plane intersecting the nucleus and using the inbuilt cell size analysis in FIJI. The staining intensity of PKC-γ was normalised to the average intensity of the dorsal horn calculated in FIJI by drawing around the dorsal column. The intensity of PKC-γ in the CST was then adjusted for area and then normalised against the fluorescence of the dorsal horn [15].

### Statistical analysis

Unless otherwise specified, two-way ANOVAs and Tukey’s post-hoc test were performed (supplementary table S1–S4). Mann Witney U test was used where the data failed a Kolmogorov-Smirnov normality test. Specific tests are detailed in the Results section. Significance was defined as p<0.05. Mean, N and S.E.M are displayed in the results tables S1 – S4. Graphs were produced in GraphPad Prism. Figures were prepared in Photoshop CS2.

## Results

### *Identification of* α-MNs and γ-MNs

α-MNs were distinguished from γ-MNs using the selective expression of NeuN and presence of C boutons on the soma [6,7], (Fig. 1A). The observed that the identified γ-MNs have a significantly smaller area and soma size compared to α-MNs in both SOD1^G93A^ and WT mice (two-way ANOVA with Tukey’s multiple comparisons test, p>0.05) (Fig. 1B-D). We found that α-MNs in SOD1^G93A^ mice are smaller compared to aged matched controls (p<0.001, Mann Whitney U Test) (Fig. 1C). There was no difference in γ-MNs soma size in SOD1^G93A^ (p>0.05, Mann Whitney U Test) (Fig 1D).

A portion of SOD1^G93A^ α-MNs were identified as “putative” α-MNs, with downregulated NeuN expression. The size of these α_p_-MNs was not significantly different from α-MNs (p>0.05, Mann Whitney U test, N = 38 α_p_-MNs, N = 22 α-MNs) and they contained C-boutons on the cell soma. Therefore, the counts were combined with α-MNs.

### Selective motoneurone loss in the cervical and lumbar spinal cord

We found extensive α-MN loss at both the cervical and lumbar levels of the spinal cord in SOD1^G93A^ mice compared to age-matched WT mice. In the cervical and lumbar spinal cords, SOD1^G93A^ mice had fewer α-MNs than WT mice at both 3- and 4-months (p<0.05) (Fig. 2B-C). We observed fewer α-MNs in the lumbar than cervical VH in SOD1^G93A^ mice, (3-Month: Lumbar 4.3 ± 0.9, Cervical 7.6 ± 0.3, 4-Month: Lumbar 4.8 ± 1.3, Cervical 5.7 ± 1.0, p<0.05) (Fig 2D/G). Suggesting, the earliest loss of motoneurones, in SOD1^G93A^ mice, is in the lumbar spinal cord.

**Figure 2:**
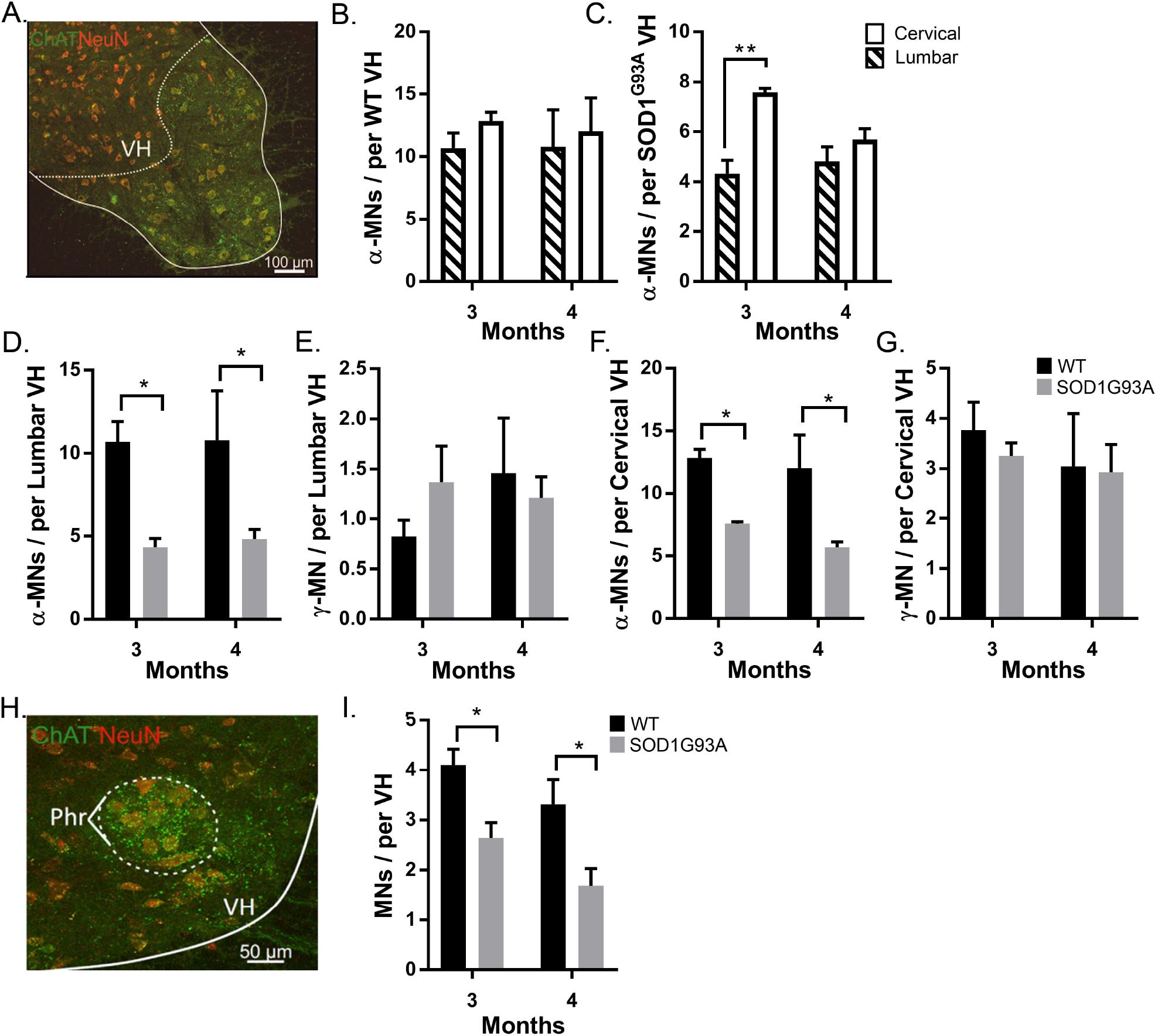
α-MN degeneration and γ-MN sparing is seen at both cervical and lumbar levels of the spinal cord of SOD1^G93A^ mice. A) Identification of the ventral horn with ChAT and NeuN. B) Average α-MN per lumbar ventral horn in 3- and 4-month WT and SOD1^G93A^ mice. C) Average α-MN per cervical ventral horn in 3- and 4-month WT and SOD1^G93A^ mice. D-E) Average α-MN per combined cervical and lumbar ventral horn in 3- and 4-month WT (D) and SOD1^G93A^ (E) mice. F) Average y -MN per lumbar ventral horn in 3- and 4-month WT and SOD1^G93A^ mice. G) Average y -MN per cervical ventral horn in 3- and 4-month WT and SOD1^G93A^ mice. H) Identification of the Phrenic nucleus by comparing ChAT and NeuN staining to the Allen mouse spinal cord atlas (http://mousespinal.brain-map.org/) I) Average MN per phrenic nerve in WT and SOD1^G93A^ mice. * denotes p<0.05** denotes p<0.01 Significance was determined by a two-way ANOVA and Tukey’s post-hoc test. Columns represent mean ± S.E.M.

There was no difference in the survival of γ-MNs between SOD1^G93A^ and WT mice at both 3- and 4-month timepoints in either spinal cord segment (p>0.05, Two-way ANOVA) (Fig. 2E/G). Hence, age, spinal segment and genotype has no effect on the survival of γ-MNs, representing the complete sparing of γ-MNs in SOD1^G93A^ mice.

### α-MNs of the phrenic nucleus experience degeneration in SOD1^G93A^ mice

As respiratory function is preserved until end stage of ALS in humans and SOD1^G93A^ mice [16] we assessed whether MN are preserved in the phrenic nucleus of SOD1^G93A^ mice. The phrenic motor pools were identified by comparing sections to the Allen mouse spinal cord atlas (http://mousespinal.brain-map.org/) (Fig. 2H). MN in the phrenic nucleus were identified as described in Materials and Methods (Fig. 1). At both 3 and 4 months, significant MN loss was observed in the phrenic motor pools of SOD1^G93A^ mice, (p<0.05 Two-way ANOVA with Tukey’s multiple comparisons test.) (Fig. 2I).

### ‘putative’ α-MNs in SOD1^G93A^ mice

A subset of α-MNs were identified as α_p_-MNs, with downregulated NeuN expression. These α_p_-MNs were included with α-MNs for analysis (Fig. 2). However, as NeuN immunoreactivity is variable in response to aging or cellular stress [17], quantification of α_p_-MNs are reported separately. The proportion of αp-MNs was no different in lumbar motor pools in SOD1^G93A^ mice compared to aged match controls (p>0.05, Two-way ANOVA).

### Integrity of the CST in the cervical and lumbar spinal cord

There was no difference in integrity of the CST axons identified by PKC-γ normalised staining intensity [15], in SOD1^G93A^ mice (p>0.05, Two-way ANOVA) (Fig. 3A-C). However, we observed a trend of decreased PKC-γ intensity on the lumbar spinal cord (p=0.052), but not cervical (p=0.38), (Two-way ANOVA with Tukey’s multiple comparisons test) (Fig. 3B-C). In 3-month SOD1^G93A^ mice, the combined CST staining intensity was 13% less than aged matched controls (Fig. 3B).

**Figure 3:**
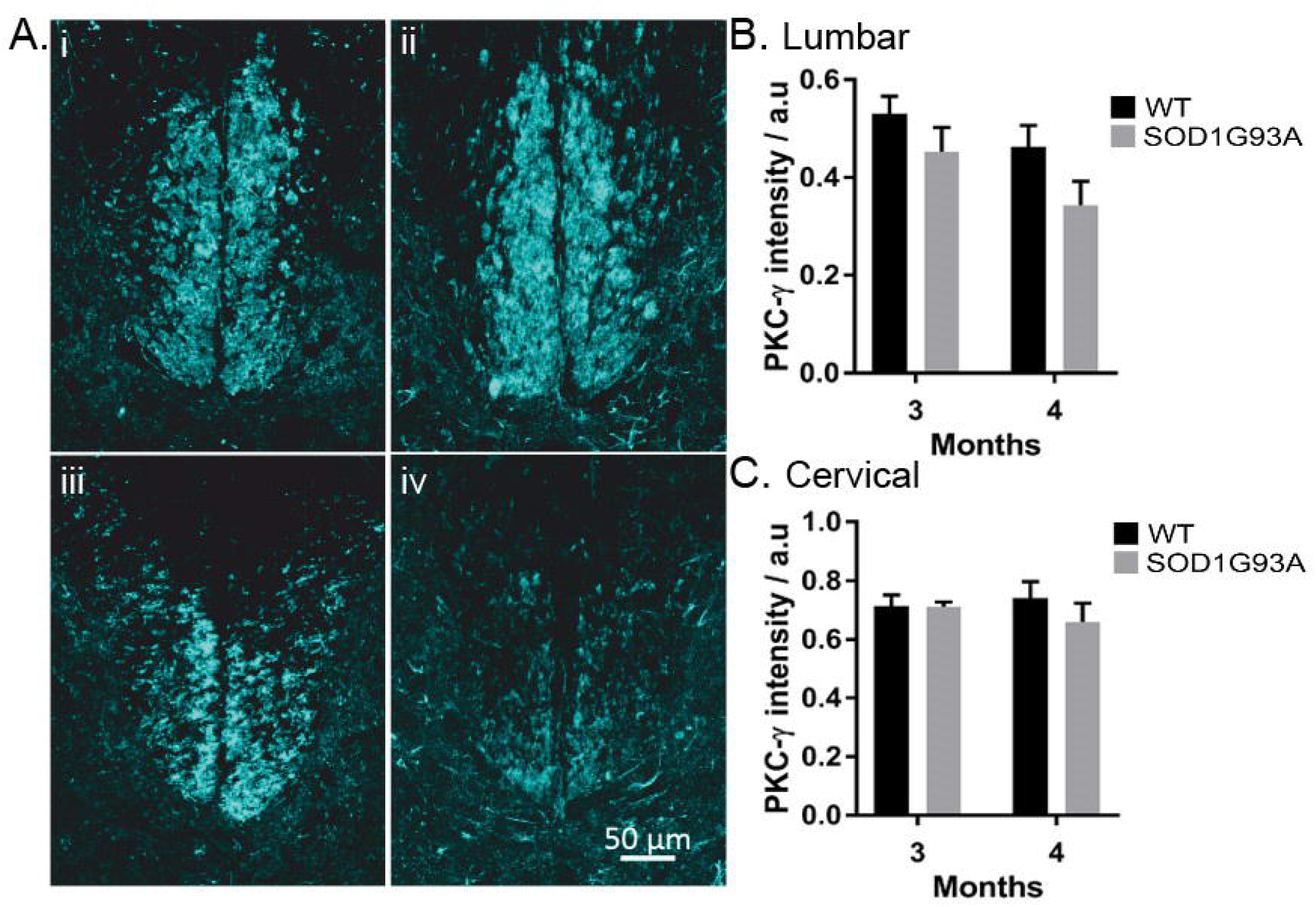
PKC-γ intensity decreases in the lumbar spinal cord of SOD1G93A mice. Ai) PKC-γ staining of a section from the 8th cervical segment in a 3-month WT mouse. Aii) PKC-γ staining of a section from the 8th cervical segment in a 4-month SOD1G93A mouse. Aiii) PKC-γ staining of a section from the 1st lumbar segment in a 3-month WT mouse. Aiv) PKC-γ staining of the 2nd lumbar segment in a 4-month SOD1G93A mouse. Scale bar reads 50 μm. B-C) Comparison of the intensity of PKC-γ staining normalised to the intensity of the dorsal horn, Lumbar (B) and Cervical (C). Columns represent mean ± S.E.M.

## Discussion

We used the selective expression of ChAT and NeuN as identifiers of α-MN over γ-MNs. Overlap in the soma sizes of α- and γ-MNs means that for accurate quantification of motor neuron survival, size analysis is insufficient, this is exacerbated further as larger, fast firing α-MNs are most vulnerable to degeneration in SOD1^G93A^ mice [5] (Fig. 1). We found that, at both cervical and lumbar levels, SOD1^G93A^ mice experience α-MN degeneration with complete γ-MN sparing (Fig. 2), similar to that reported previously [4]. We further identified MN loss in the phrenic motorpools in SOD1^G93A^ mice (Fig. 2). The integrity of the CST followed a similar caudal to rostral degeneration in the cervical and lumbar spinal cord (Fig. 3). Our results suggest that MN degeneration follows a caudal to rostral progression, with complete sparing of γ-MNs in the SOD1^G93A^ mouse model of ALS.

By 3 months, there was degeneration of α-MNs with complete sparing of γ-MNs, in both the cervical and lumbar spinal cord (Fig 2.). The observed sparing of γ-MNs is supported by previous studies reporting a decrease in α-MNs in SOD1^G93A^ mice using retrograde labelling [18]. Sparing of γ-MNs was also reported in TDP-43 and FUS models of ALS [4], suggesting it is a common feature of ALS pathology. Interestingly, the survival of γ-MNs may further exacerbate excitotoxicity in the remaining α-MNs, mediated through increased proprioceptive feedback from Ia afferents to compensate for the loss of α-MNs to produce the required force. This hypothesis is supported by the absence of degeneration in the oculomotor nerve and Onuf’s nucleus, which lack proprioceptive Ia afferent feedback from the muscle spindle [4].

We found that there were fewer α-MNs in the lumbar compared to cervical motor pools. This suggests that cervical α-MNs are spared until a later stage in the disease (Fig. 2). Surviving α-MNs in the cervical spinal cord could be the foundation for preserved forelimb function until disease end-stage [2]. One possibility is that the spinal motor circuit compensates for α-MNs loss, through reorganisation of the surviving motor circuit, such as Renshaw and cholinergic rewiring [19,20] or afferent sprouting in absence of the CST [15,21].

A similar progression of α-MN loss, preceding motor impairment, was observed in the phrenic nucleus (Fig. 2). In SOD1^G93A^ mice, breathing capacity (tidal volume and minute pulmonary ventilation) is preserved until drastic declines in the final two days of life [22]. This is despite the extensive α-MN degeneration we identified (Fig. 2). Two factors may influence the absence of respiratory motor impairment, despite MN loss. First, the relative lack of γ-MNs, reducing the potential for proprioceptive feedback to exacerbate degeneration. Second, the difference in descending cortico-spinal pathways, as the phrenic nucleus is instead innervated by the bulbospinal tracts. Even though, bulbospinal tracts undergo degeneration in SOD1^G93A^ model of ALS [23], degeneration of the tract may not exacerbate α-MN loss through afferent sprouting, as observed by damage to the CST [15].

At 4 months we observed a trend of decreased PKC-γ intensity on the lumbar spinal cord (p=0.052), but not cervical (p=0.38). This preferential degeneration of the lumbar CST has been previously reported in SOD1^G93A^ mice at 120 days of age [23,24]. Suggesting that, in line with corticofugal synaptopathy model, supraspinal motoneurones targeting the lumbar spinal cord are more susceptible to degeneration [9]. Loss of the CST has previously been shown to induce proprioceptive afferent sprouting in the adult spinal cord, alongside an increase in glutamatergic boutons connected to spinal motor neurons [15]. If similar plasticity mechanisms are evoked in the lumbar segment of SOD1^G93A^ mice, this could further exacerbate glutamate excitotoxicity of α-MNs. Our findings agree with previous reports but in the same animal, suggesting that the morbidity in ALS is the effect of the sum of its parts. All the events leading to cell death, along with the plastic changes induced by loss of pathways and neurones is what exacerbates the system leading it towards its diminishing ability to maintain activity.

## Conclusion

In summary, there is selective degeneration of α-MNs and complete sparing of γ-MNs at both lumbar and cervical spinal segments in SOD1^G93A^ mice. The α-MNs lumbar spinal cord appear more susceptible to degeneration than the cervical spinal cord at 3 months in SOD1^G93A^ mice. This caudal to rostral progression is further observed in the integrity of the CST. These results raise the question of how the sparing of γ-MNs and CST degeneration could further exacerbate α-MNs loss and motor impairment in ALS.

## Acknowledgements

The authors would like to thank Celine de Kamps with image analysis and acknowledge funding from The Faculty of biological sciences, BMSC3301 programme for undergraduate projects, BIOL5294 program for MSc Bioscience research projects.

## Author contribution

AJK, TP performed the experiments, analysed the data. AJK, TP, SC, RM designed the experiments and wrote the manuscript.

## Data Accessibility

Data are available on request from the corresponding author.

---

**Figure S1:**
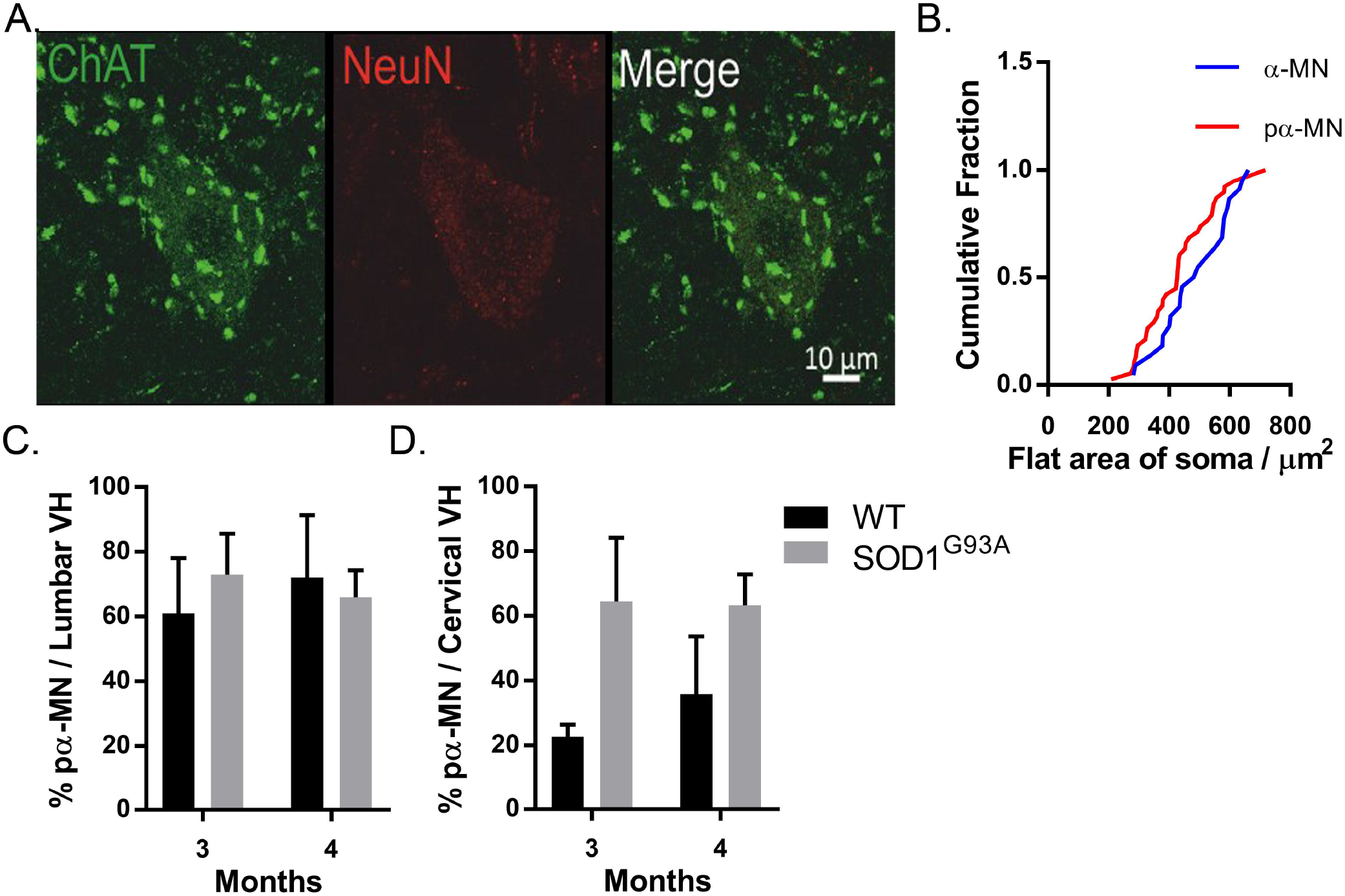
*Identification of* α_p_-MNs

**Table S1:** Size comparison of α-MNs and γ-MNs between SOD1^G93A^ and WT mice:

| Group | Statistical test | N | P Values |
| --- | --- | --- | --- |
| SOD1<sup>G93A</sup> v WT<br>(α-MNs v γ-MNs) | Two-Way ANOVA,<br>Tukey's multiple comparison test | SOD1<sup>G93A</sup>: α-MNs (4) γ-MNs (4)<br>WT: α-MNs (4) γ-MNs (4) | Genotype: 0.0015<br>MN classification: <0.001<br>Interaction: 0.0087 |
| Distribution analysis:<br>α-MNs: SOD1<sup>G93A</sup> v WT | Mann Whitney U Test. | SOD1<sup>G93A</sup>: α-MNs (60)<br>WT: α-MNs (60) | P <0.001 (Two tailed) |
| Distribution analysis:<br>γ-MNs: SOD1<sup>G93A</sup> v WT | Mann Whitney U Test. | SOD1<sup>G93A</sup>: γ-MNs (4)<br>WT: γ-MNs (4) | P <0.001 (Two tailed) |
| Distribution analysis:<br>$\alpha_p$-MNs v WT α-MNs | Mann Whitney U Test. | $\alpha_p$-MNs (22)<br>WT: α-MNs (38) | P 0.912 (Two tailed) |

**Table S2:** MNs loss in between 3- and 4- months in SOD1^G93A^ and WT mice: (* Indicates significance.)

| Group | Statistical test | N | P Values |
| --- | --- | --- | --- |
| **α-MNs Lumbar**<br>SOD1<sup>G93A</sup> v WT<br>(3-Months v 4-Months) | Two-Way ANOVA,<br>Tukey's multiple comparison test | 3-Month: WT (5), SOD1<sup>G93A</sup> (3),<br>4-Month: WT (3), SOD1<sup>G93A</sup> (5). | *Genotype: 0.0009<br>Age: 0.8368<br>Interaction: 0.8929 |
| **γ-MNs Lumbar**<br>SOD1<sup>G93A</sup> v WT<br>(3-Months v 4-Months) | Two-Way ANOVA,<br>Tukey's multiple comparison test | 3-Month: WT (5), SOD1<sup>G93A</sup> (3),<br>4-Month: WT (3), SOD1<sup>G93A</sup> (5). | Genotype: 0.508<br>Age: 0.705<br>Interaction: 0.07 |
| **α-MNs Cervical**<br>SOD1<sup>G93A</sup> v WT<br>(3-Months v 4-Months) | Two-Way ANOVA,<br>Tukey's multiple comparison test | 3-Month: WT (5), SOD1<sup>G93A</sup> (3),<br>4-Month: WT (3), SOD1<sup>G93A</sup> (5). | *Genotype: 0.0002<br>Age: 0.2472<br>Interaction: 0.6382 |
| **γ-MNs Cervical**<br>SOD1<sup>G93A</sup> v WT<br>(3-Months v 4-Months) | Two-Way ANOVA,<br>Tukey's multiple comparison test | 3-Month: WT (5), SOD1<sup>G93A</sup> (3),<br>4-Month: WT (3), SOD1<sup>G93A</sup> (5). | Genotype: 0.9457<br>Age: 0.5898<br>Interaction: 0.7307 |
| **WT α-MNs** | Two-Way ANOVA, | 3-Month: Lumbar (5), Cervical (5), | Spinal level: 0.2139 |
| Lumbar v Cervical<br>(3-Months v 4-Months) | Tukey's multiple comparison test | 4-Month: Lumbar (3), Cervical (3). | Age: 0.3193<br>Interaction: 0.9633 |
| **SOD1<sup>G93A</sup> α-MNs**<br>Lumbar v Cervical<br>(3-Months v 4-Months) | Two-Way ANOVA,<br>Tukey's multiple comparison test | 3-Month: Lumbar (5), Cervical (5),<br>4-Month: Lumbar (3), Cervical (3). | *Spinal level: 0.002<br>Age: 0.205<br>*Interaction: 0.0415 |
| **WT γ-MNs**<br>Lumbar v Cervical<br>(3-Months v 4-Months) | Two-Way ANOVA,<br>Tukey's multiple comparison test | 3-Month: Lumbar (5), Cervical (5),<br>4-Month: Lumbar (3), Cervical (3). | Spinal level: 0.0022<br>Age: 0.9420<br>Interaction: 0.2664 |
| **SOD1<sup>G93A</sup> γ-MNs**<br>Lumbar v Cervical<br>(3-Months v 4-Months) | Two-Way ANOVA,<br>Tukey's multiple comparison test | 3-Month: Lumbar (5), Cervical (5),<br>4-Month: Lumbar (3), Cervical (3). | Spinal level: 0.2764<br>Age: 0.4078<br>Interaction: 0.2807 |

**Table S3:** α-MNs counts in the phrenic nucleus between 3 and 4 months

| Group | Statistical test | N | P Values |
| --- | --- | --- | --- |
| **α-MNs Phrenic nucleus**<br>SOD1<sup>G93A</sup> v WT<br>3-Months v 4-Months | Two-Way ANOVA,<br>Tukey's multiple comparison test | 3-Month: WT (5), SOD1<sup>G93A</sup> (3).<br>4-Month: WT (3), SOD1<sup>G93A</sup> (5). | *Genotype: 0.001<br>Age: 0.0312<br>Interaction: 0.8269 |

**Table S4:** PKC-y intensity between 3 and 4 months in SOD1 and WT mice:

| Group | Statistical test | N | P Values |
| --- | --- | --- | --- |
| **Lumbar**<br>SOD1<sup>G93A</sup> v WT<br>(3-Months v 4-Months) | Two-Way ANOVA,<br>Tukey's multiple comparison test | 3-Month: WT (5), SOD1<sup>G93A</sup> (4),<br>4-Month: WT (3), SOD1<sup>G93A</sup> (3). | Genotype: 0.0521<br>Age: 0.0786<br>Interaction: 0.6485 |
| **Cervical**<br>SOD1<sup>G93A</sup> v WT<br>(3-Months v 4-Months) | Two-Way ANOVA,<br>Tukey's multiple comparison test | 3-Month: WT (5), SOD1<sup>G93A</sup> (3),<br>4-Month: WT (3), SOD1<sup>G93A</sup> (3). | Genotype: 0.3866<br>Age: 0.7955<br>Interaction: 0.4244 |

